# Predictive Modeling of Immune Escape and Antigenic Grouping of SARS-CoV-2 Variants

**DOI:** 10.1101/2025.05.28.656328

**Authors:** Arshan Nasir, Diana Lee, Laura E. Avena, Daniela Montes Berrueta, Tessa Speidel, Kai Wu, Yadunanda Budigi, Andrea Carfi, Guillaume B. E. Stewart-Jones, Darin Edwards

## Abstract

The ongoing adaptive evolution of SARS-CoV-2 is characterized by the continued emergence of novel variants that escape from previously acquired infection- and/or vaccination-derived immunity. This continued SARS-CoV-2 variant evolution has necessitated annual vaccine updates to better match circulating viral variants. To optimize protection against emerging variants of interest and concern, a reliable means of predicting the immune escape of novel variants is needed to enable at-risk preparation of new vaccines. Herein, we describe the development and applications of a risk calculator that uses statistical modeling to predict the immune escape of emerging variants. The calculator utilizes previously published spike-antibody epitope and escape profiles and *in vitro* neutralization assessment of a large panel of pseudotyped SARS-CoV-2 variants evaluated against clinical sera. The calculator enables the grouping of antigenically related SARS-CoV-2 variants to guide strain selection for at-risk vaccine design and preparation, in anticipation of potential future requests by the global public health agencies. Here, we demonstrated the strain selection exercises for the XBB.1.5- and JN.1/KP.2-adapted mRNA-1273 COVID-19 vaccines in the 2023-2024 and 2024-2025 seasons, respectively, which were supported by both the risk calculator and preclinical and clinical immunogenicity data and were later recommended by the global public health agencies.

## Background

Since the emergence of SARS-CoV-2 in 2019, the virus has shown a remarkable capacity for rapid antigenic evolution, giving rise to novel variants with distinct immune evasion and transmission properties^1^. These immune-evasive variants often carry specific mutations in key regions of the viral genome such as the surface spike glycoprotein (S), which plays a crucial role in viral entry into the host cells^1, 2^. Mutations in S, especially in the receptor-binding domain (RBD), can affect viral infectivity, transmissibility, and immune recognition, potentially leading to altered disease outcomes and immune escape^1, 3^. Timely identification and characterization of emerging high-risk immune-evasive SARS-CoV-2 variants is thus vital for implementing effective public health response measures, including targeted testing and proactive adjustments to approved vaccines to optimize continued efficacy against evolving strains and to mitigate their risk to global public health^4^.

In this manuscript, we describe the development and application of a risk calculator that analyzes new mutations observed in the RBD of new SARS-CoV-2 variants and uses statistical modeling to predict the impact on neutralization by post-vaccination serum. The risk calculator provides a quick and quantitative *in silico* assessment of the potential immune escape of emerging variants, enabling grouping of antigenically related strains and prioritization of strains for at-risk vaccine development including assessment of preclinical and clinical immunogenicity. Here we demonstrate the utility of this strategy in at-risk strain selection and preparation of updated mRNA-1273 COVID-19 vaccines in the 2023-24 and 2024-25 season.

## Methods

### Clinical trial and serum neutralization activity

Serum neutralization activity was measured against different SARS-CoV-2 variants in a pseudovirus (PsV)-based model to train the risk calculator. Specifically, Day 29 sera (4 weeks after the fourth dose) from healthy adult participants (n = 20) enrolled in the phase 2/3 Part H study (NCT04927065; the immunization history of the study cohort is reported in **Table 1**) were collected and assessed against each of the 18 pseudotyped variants for neutralization activity (the pseudoviruses assessed are listed in **Table 2**)^5, 6^. Study protocols and results were reported previously^5, 6^. Eligible participants received a primary series (2 doses) of mRNA-1273 (the original vaccine targeting the ancestral Wuhan-Hu-1 strain), followed by a third booster dose of mRNA-1273, and a fourth booster dose of mRNA-1273.222 (the 2022-2023 bivalent vaccine formulation targeting the ancestral Wuhan-Hu-1 and BA.4/BA.5 strains)^5, 6^. Among the 20 participants, 10 had a history of prior infection. This study was conducted in accordance with the Good Clinical Practice guidelines of the International Council Harmonisation of Technical Requirements for Pharmaceuticals for Human Use, and applicable local regulations. The central institutional review board approved the protocol and the consent forms. All participants provided written informed consent.

**Table 1.** Immunization history of the participants in the phase 2/3 Part H study cohort.

| **Trial** | **Cohort** | **Doses 1 and 2**<br>**(primary series)** | **Dose 3 (booster)** | **Dose 4 (booster)** | **Time point** | **Sample size** |
| --- | --- | --- | --- | --- | --- | --- |
| Phase 2/3<br>NCT04927065 | Part H | mRNA-1273 100 µg | mRNA-1273 50 µg | mRNA-1273.222 50 µg | Day 29<br>post-dose 4 | N = 20 |

**Table 2.**
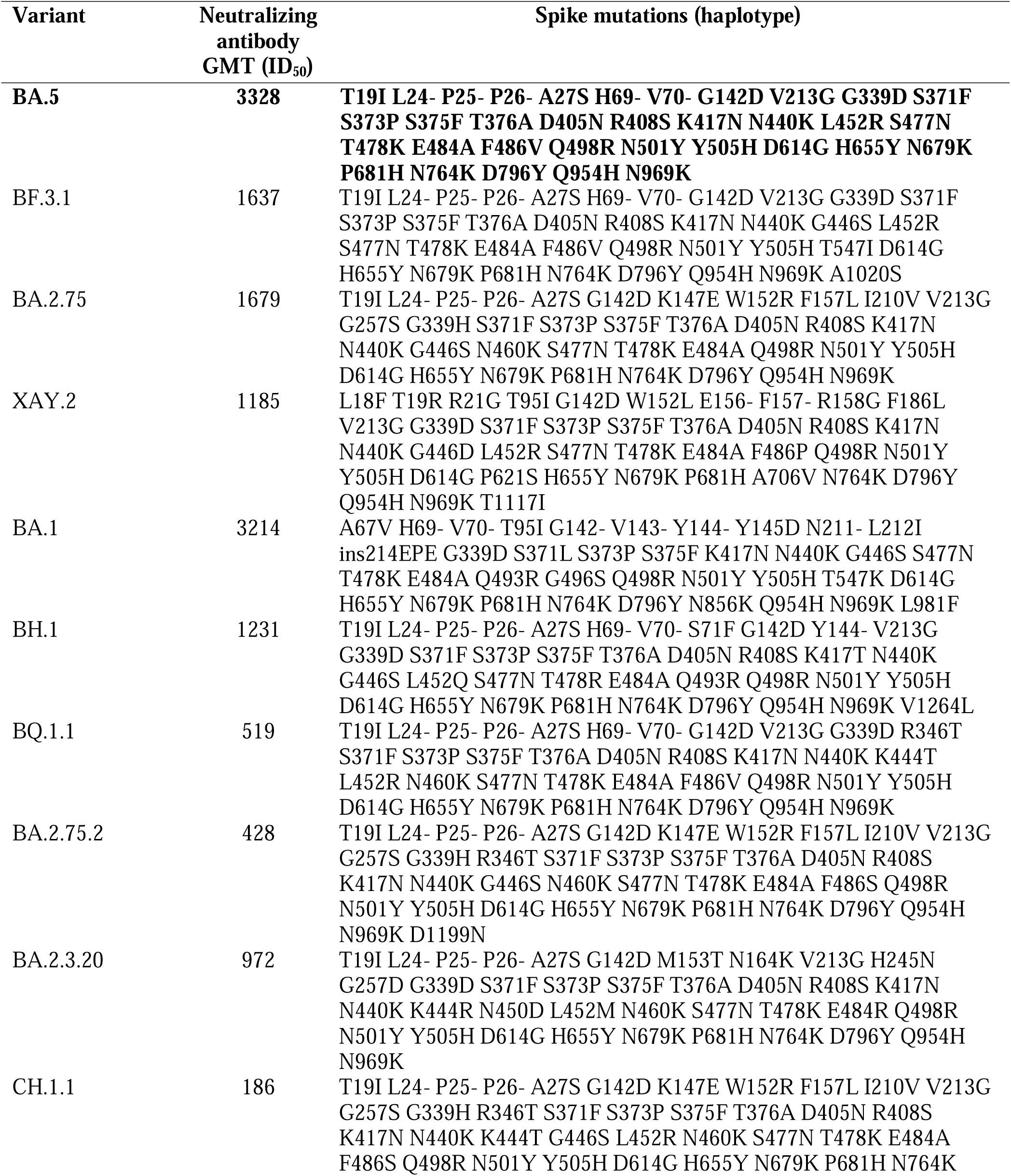
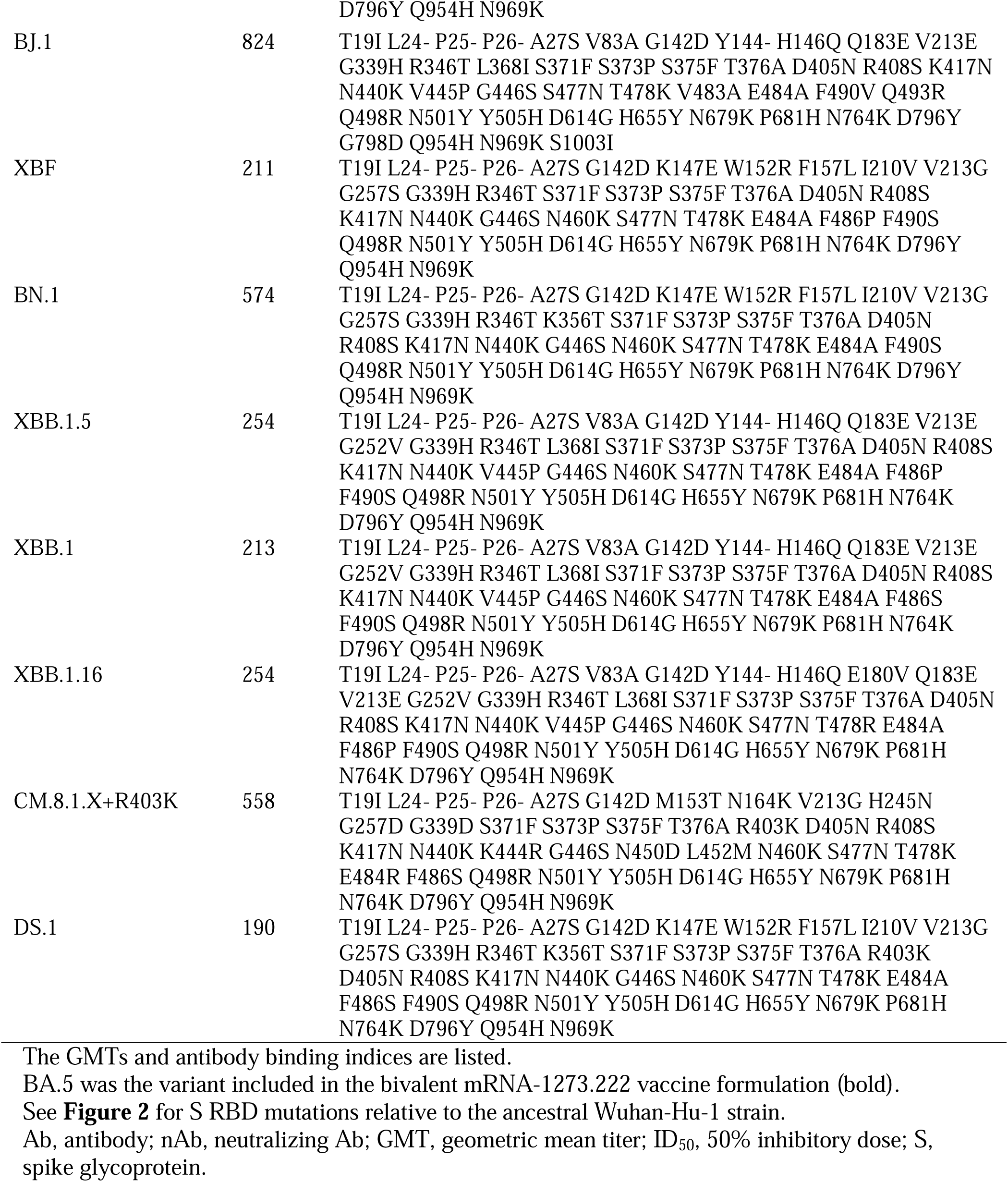
Neutralization data for pseudotyped SARS-CoV-2 variants against sera from the mRNA-1273.222 phase 2/3 Part H clinical trial.

### Antibody binding data and escape profiles

Antibody S epitopes and antibody escape profiles were retrieved from previously published datasets from the research groups of Yunlong Cao and Jesse Bloom^7–9^. These datasets describe antibody and S contact positions for >4,000 monoclonal antibodies determined via deep-mutation scanning, along with individual mutations and sites likely to cause loss of antibody binding (and by extension, immune escape)^7–9^. From these datasets, we calculated an antibody binding index that describes the potential immune escape for a S variant by calculating the fraction of antibodies likely to either completely or partially escape due to detected mutations in the S variant (similar to the index published previously by Jesse Bloom’s group)^10^. The antibody binding index thus aggregates escape over mutated sites and is bounded between 0 and 1, with 1 indicating complete escape from all antibodies in the dataset.

### Predictive modeling of immune escape

A linear model that uses antibody binding index (predictor) to predict loss in neutralization (dependent variable) was developed. The current version of the model was trained on the neutralization data from the phase 2/3 Part H study (NCT04927065; **Table 1**). The same model was used to extrapolate neutralization scores for newer variants against the more recent vaccines, including mRNA-1273.815 (the 2023-2024 monovalent vaccine formulation targeting the XBB.1.5 variant), mRNA-1273.167 (the 2024-2025 monovalent vaccine formulation targeting the JN.1 variant), and mRNA-1273.712 (the 2024-2025 monovalent vaccine formulation targeting the KP.2 variant). Of note, a previous iteration of the model trained on the mRNA-1273 phase 1 data (NCT04283461) was built to forecast immune escape from exposure to the Wuhan-Hu-1 strain (**Fig. S1**) but is no longer under active development. For the latest vaccines, the risk scores account for prior immune exposures by ignoring mutations that have previously appeared in any of the previous vaccine variants (eg, BA.4/BA.5, XBB.1.5, JN.1, or KP.2). Hence, the calculator attempts to mimic the complex immunization history in the human population acquired via a series of immunizations by assuming complete (and lasting) immunity to mutations already experienced by the human population.

### Recombinant vesicular stomatitis virus (VSV)-PsV assay

Codon-optimized full-length S genes for the 18 pseudotyped variants (**Table 2**) were cloned into a pCAGGS vector^11, 12^. The S gene mutations are listed in **Table 2**. To generate VSVΔG-based SARS-CoV-2 PsV, BHK-21/WI-2 cells were transfected with the S expression plasmid and infected by VSVΔG-firefly-luciferase as described previously^11–13^. Vero E6 cells (ATCC, CRL-1586) were used as target cells for the neutralization assay and were maintained in Dulbecco’s Modified Eagle Medium (DMEM) supplemented with 10% fetal bovine serum (FBS). To perform the neutralization assay, human serum samples were heat-inactivated for 30 minutes at 56°C, and serial dilutions were made in DMEM supplemented with 10% FBS. The diluted serum samples or culture medium (serving as a virus-only control) were mixed with VSVΔG-based SARS-CoV-2 PsV and were incubated at 37°C for 45 minutes. The inoculum virus or virus– serum mix was subsequently used to infect Vero E6 cells for 18 hours at 37°C. At 18 hours after infection, an equal volume of One-Glo reagent (Promega, E6120) was added to the culture medium for readout using a BMG PHERastar-FSX plate reader. The percentage of neutralization was calculated based on relative light units of the virus control and was subsequently analyzed using a four-parameter logistic curve (GraphPad Prism version 10.2.1).

### Sequence analysis

Newly published SARS-CoV-2 genetic sequences from the Global Initiative on Sharing All Influenza Data (GISAID) database^14^ were downloaded and the Nextclade algorithm (current version 3.8.2)^15^ was used for viral genome alignment, quality control checks, mutation calling, and clade assignment. S mutations were called relative to the ancestral Wuhan-Hu-1 strain (GenBank Accession ID: MN908947) using in-house Python (version 3.12.2) scripts. Filtered S variants were categorized using the assigned risk score. Variants that scored >2 against the most recently authorized vaccines were prioritized for preclinical evaluation and assessment or for continued monitoring of epidemiologic growth.

### Statistical analysis

Model significance was evaluated using an *F* statistic^16^. Model assumptions were validated formally using the Shapiro-Wilk and Breusch-Pagan tests for normality and equal variances, respectively^17, 18^, and visually by plotting the distribution of the fitted values versus the residuals. The input to the risk assessment model was the S amino acid sequence (or a list of RBD mutations) for a given SARS-CoV-2 variant. The output was a numerical value (called risk score) that described the expected drop in neutralization for the variant from previously approved vaccines.

### Phylogenetic analysis

The S sequences for major SARS-CoV-2 variants of interest were extracted via the Nextclade pipeline. The S alignment was used as input to the IQTREE program (version 2.3.5) for a reconstruction of maximum-likelihood phylogeny^19^. We ran 10 independent tree searches, stopping after 100 iterations in each search, and used the HIVw+F+I+G4 substitution model (suggested by the ModelFinder program) for all searches.^20^. A phylogenetic tree with the best likelihood was selected and rooted *a posteriori* in the branch leading to the ancestral (Wuhan-Hu-1) strain. The tree was visualized via FigTree (version 1.4.4)^21^.

## Results

### Quantitative assessment of immune escape

The risk assessment process includes weekly analysis of new SARS-CoV-2 genetic data from the public repositories (eg, the GISAID database and GenBank)^14^ and the use of a statistical modeling strategy to assign a strain-level risk score to the emerging variants, which informs at-risk preparation of vaccine constructs for variants with the potential to substantially escape preexisting immunity in the human population. A key goal of the process is to predict immune escape, based on an analysis of amino acid changes in the S variant, against a background of preexisting immunity from specific vaccines. To inform and train the calculator, we used antibody S binding data for >4,000 monoclonal antibodies from published studies^7–9^ and human serum neutralization activity that we measured against a panel of SARS-CoV-2 variants using a pseudovirus neutralization assay (**Table 2**). The data were processed to fit a linear regression model that predicts loss in neutralization (dependent variable) as a function of antibody binding index (predictor), defined by the fraction of antibodies that may have lost binding capabilities due to observed mutations (similar to that described in Greaney AJ, et al)^10^. The statistical model is described by the following equation:

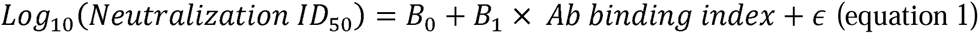

Where *β*_0_ (intercept) is the expected neutralization when antibody binding index = 0 (ie, the 50% inhibitory dose [ID_50_] neutralization titer for the matched variant included in the vaccine). *Β*_1_ (slope) is the (exponential) decrease in neutralization for a S variant per 1% increase in the antibody binding index and *ε* is the error term or residuals. Errors were assumed to be normally and independently distributed (*W* statistic, 0.95; *p* = 0.36, Shapiro-Wilk test) with equal variances (Lagrange multiplier statistic, 0.69; *p* = 0.41, Breusch-Pagan test).

### Antibody binding index correlates strongly with the experimental measurements of immune escape

The antibody binding index was a significant predictor of neutralization drop (*R*^2^ = 0.7; *F* value (1, 16) = 38.05; *p* < 0.01) when tested on 18 different SARS-CoV-2 variants that included major variants that had become epidemiologically relevant up until April 30, 2023, the last cutoff date for the latest experimentally validated and trained model (**Fig. 1**, **Table 2**). For example, tested variants included BA.2 descendants, such as members from the BA.2.75 sublineage, which are characterized by a distinct N-terminal domain (NTD) mutation profile; BA.2.3.20 and its descendants, which are characterized by unique S substitutions not seen previously; BA.5 and its descendants (BQ.1.1), which became predominant in 2022 and prompted a vaccine update; and several recombinant strains such as the well-characterized XBB family (XBB.1, XBB.1.5, and XBB.1.16), which prompted a vaccine update in the 2023-2024 season (**Fig. 2**). The estimated regression function, *Log*_0_(*Neutralization ID*_50_) = 3.47 - 3.15 x *Ab binding index*, suggests that neutralization titers decrease by ∼7.5% in the original scale for each unit (1%) increase in the antibody binding index.

**Fig. 1.**
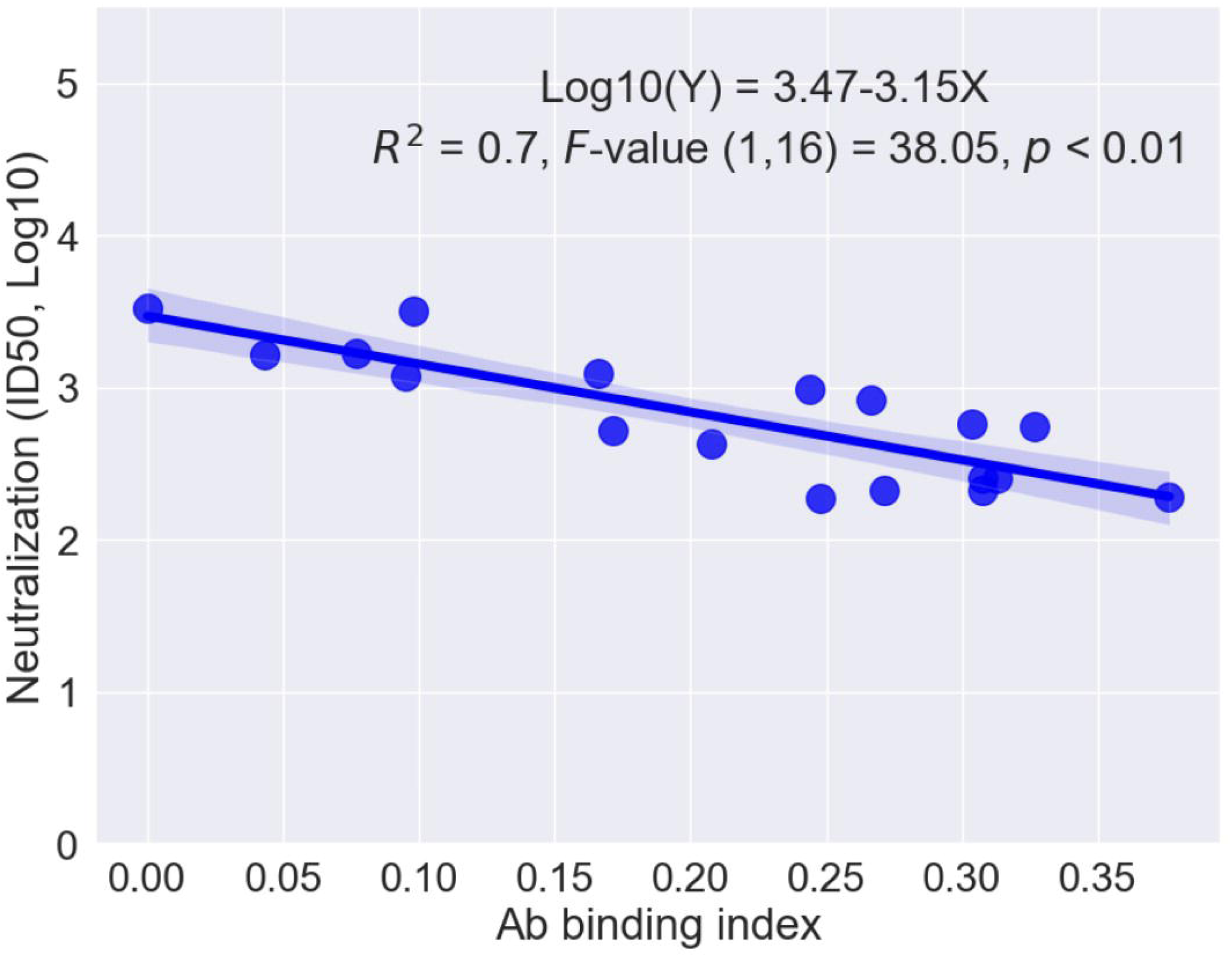
Statistical relationship between antibody binding index (*x*-axis) and serum neutralization for SARS-CoV-2 variants evaluated against clinical sera from the mRNA-1273.222 phase 2/3 Part H study (*y*-axis). The estimated regression function is listed, along with the statistical test for significance. Ab, antibody; ID_50_, 50% inhibitory dose.

**Fig. 2.**
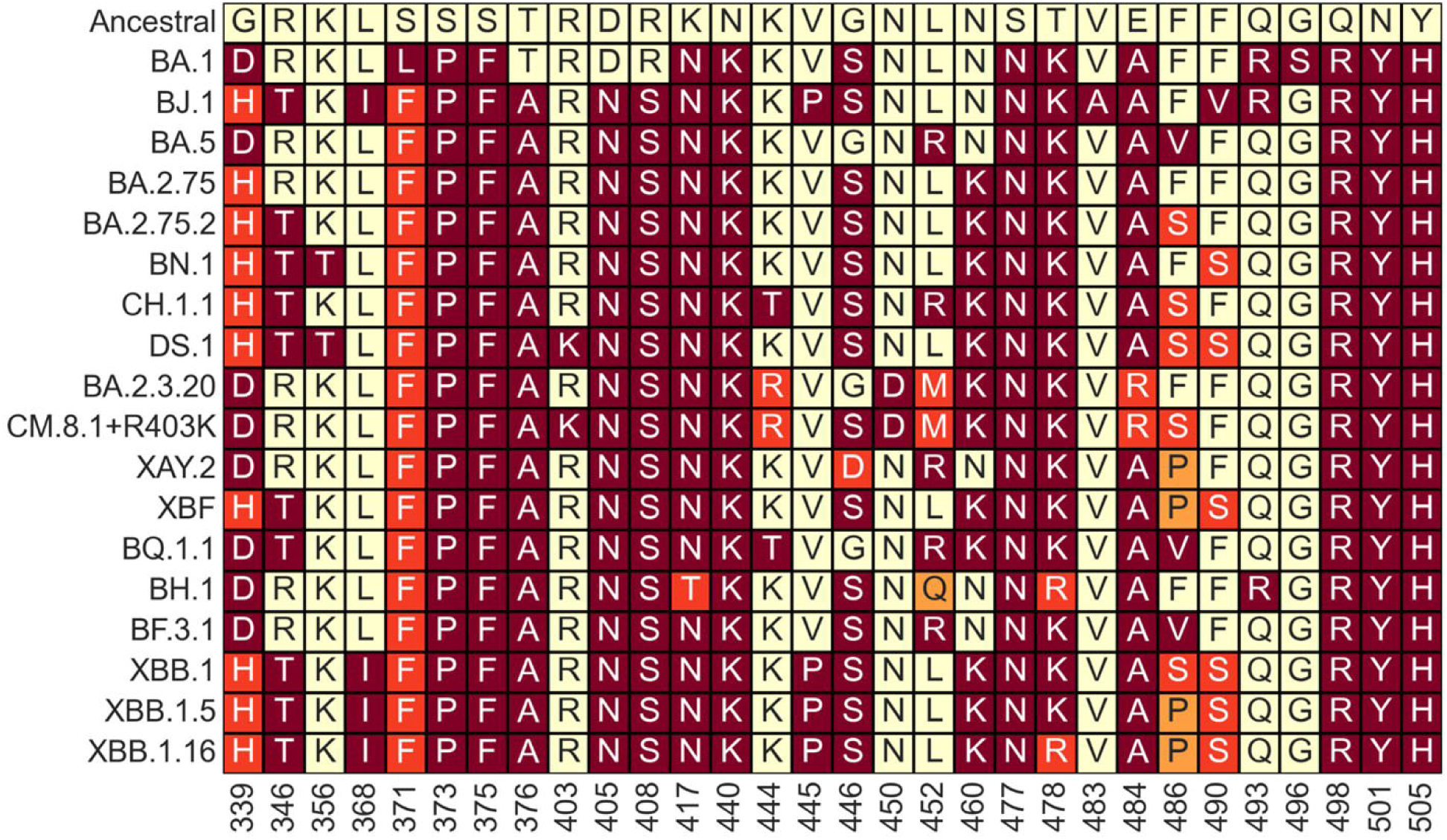
Antigenic diversity of variants assessed against clinical sera. S RBD mutations in pseudotyped SARS-CoV-2 variants that were evaluated against sera from the mRNA-1273.222 phase 2/3 Part H study. See Table 2 for full list of S mutations. The *x*-axis indicates the amino acid position relative to the ancestral strain (Wuhan-Hu-1). Mutations are indicated in different colors. Some sites have mutated more than once, such as sites 339, 371, 417, 444, 446, 452, 478, 486, and 490. RBD, receptor-binding domain; S, spike glycoprotein.

### Risk calculator enables antigenic grouping of variants and predicts cross-neutralization from approved vaccines

A key utility of the risk calculator is its potential to calculate risk scores for emerging variants against licensed SARS-CoV-2 vaccine compositions (**Fig. 3**), as well as new variant vaccine candidates under development. The risk scoring and ranking enables rapid identification and prioritization of immune-evasive variants based solely on *in silico* analysis for further epidemiologic monitoring and potential commercial development, if needed. The predicted risk scores for a range of variants against a panel of previously or currently authorized/approved vaccines, including mRNA-1273 (the original monovalent vaccine formulation targeting the ancestral Wuhan-Hu-1 strain), mRNA-1273.222 (the 2022-2023 bivalent vaccine formulation targeting the Wuhan-Hu-1 and BA.5 strains), mRNA-1273.815 (the 2023-2024 monovalent vaccine formulation targeting the XBB.1.5 strain), mRNA-1273.167 (the 2024-2025 monovalent vaccine formulation targeting the JN.1 strain), and mRNA-1273.712 (the 2024-2025 monovalent vaccine formulation targeting the KP.2 strain for use in the United States and Canada) vaccines are shown in **Fig. 3**. The RBD mutations in the major SARS-CoV-2 variants that became epidemiologically relevant in the 2023-2024 and 2024-2025 seasons are shown in **Fig. 3a**, along with few historically successful variants, and the predicted risk scores from approved SARS-CoV-2 vaccines are shown in **Fig. 3b**. In the 2023-2024 season, the risk calculator indicated that XBB.1.5 had a greater risk score (9.29) against the mRNA-1273.222 vaccine authorized at that time, compared with other co-circulating strains such as BQ.1.1, BA.2.75.2, CH.1.1, and BA.2.3.20, which had substantially lower risk scores. The higher risk score associated with XBB.1.5 was likely due to the presence of several new and convergent RBD mutations, including G339H, R346T, L368I, V445P, G446S, N460K, F486P, and F490S (**Fig. 3a**) that conferred a higher degree of immune escape to XBB.1.5 compared with other strains circulating at that time. The risk calculator also enables antigenic grouping of variants. For example, XBB-derived sublineages XBB.1.16, XBB.2.3.2, EG.5.1, FL.1.5.1, HV.1, and HK.3.1, which emerged either during the XBB.1.5 infection wave or soon after, had comparable risk scores to XBB.1.5 (1- to 2.25-fold difference). Collectively this information derived from the risk analysis suggested that an XBB.1.5 vaccine was likely to neutralize all those strains and was likely the most suitable candidate for a vaccine update (**Fig. 3b**).

**Fig. 3.**
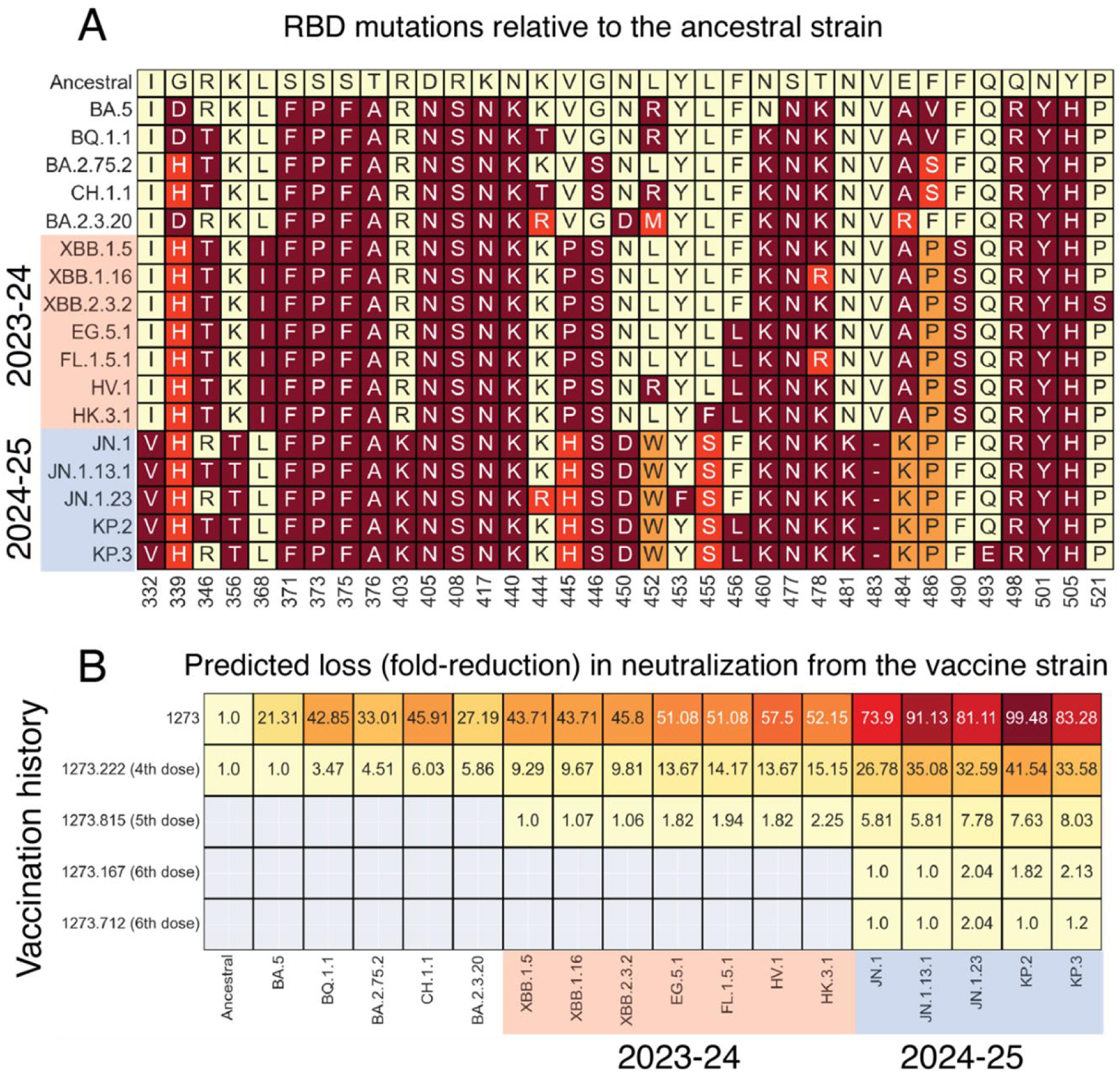
Antigenic grouping of variants in the 2023-2024 and 2024-2025 seasons and predicted immune escape by vaccination history. (A) S RBD mutations in SARS-CoV-2 variants that were epidemiologically relevant in the 2023-2024 and 2024-2025 seasons. The *x*-axis indicates the amino acid position relative to the ancestral (Wuhan-Hu-1) strain. Mutations are indicated in different colors. Some sites have mutated have more than once, such as 339, 444, 445, 452, 455, 478, 484, and 486. (B) The matrix lists the predicted immune escape for the same variants against different immunization backgrounds. The values inside the matrix are the predicted fold-drop in the ID_50_ neutralizing antibody GMTs on the vaccine strain, derived from the statistical model described in Figure 1. GMT, geometric mean titer; ID_50_, 50% inhibitory dose; RBD, receptor-binding domain; S, spike glycoprotein.

In late 2023, JN.1 (a descendant of BA.2.86 lineage) emerged and quickly displaced the XBB subvariants as the predominant circulating strain globally by early 2024. JN.1 was characterized by ∼60 S mutations relative to the ancestral Wuhan-Hu-1 strain and >30 mutations compared with the XBB.1.5 strain,^22^ which was included in the mRNA-1273.815 vaccine for the 2023-2024 season. Some notable mutations in the JN.1 S included K356T, which introduced a new N-linked glycosylation site in the RBD; N460K, which likely improved ACE2 binding affinity; L455S, which likely improved the transmission capabilities of JN.1; and several additional NTD and RBD substitutions, insertions, and deletions likely to improve immune escape^23–25^. Initial testing revealed a 5.8-fold-drop in nAb titers to JN.1 compared with the XBB.1.5 strain in the sera from mRNA-1273.815-boosted individuals^11^. We used these data for predictive modeling of JN.1 and its subvariants that had emerged and/or were co-circulating at the time of strain selection, including JN.1.13.1, JN.1.23, KP.2, and KP.3. All JN.1 subvariants had comparable risk scores to JN.1 and KP.2 (∼1-2-fold-difference), suggesting that initial JN.1 and KP.2 were well suited as candidate vaccine strains for continued development candidates.

### mRNA-1273.167 (JN.1) and mRNA-1273.712 (KP.2) vaccines are likely to cross-neutralize currently emerging JN.1 subvariants

At the time of this manuscript writing (January 2025), XEC (a recombinant strain with a S breakpoint) has become dominant globally, replacing KP.3.1.1 (a JN.1 subvariant), the previous dominant strain. The XEC variant is a recombinant virus involving recombination between 2 JN.1 subvariants (KP.3.3 and KS.1.1)^26^. The XEC S RBD is identical to the KP.3.1.1 RBD (**Fig. 4a**) but has a different NTD glycosylation site compared with KP.3.1.1. (T22N in XEC vs. S31-in KP.3.1.1); XEC is currently the most frequently reported variant both in Europe and North America but has been trending down in recent weeks^27^. In addition, there are some geographical differences in the distribution of SARS-CoV-2 variants in some countries and new upcoming challengers. For example, NB.1 is dominant in samples from China^28–30^. NB.1 has evolved from XDV, a complex recombinant virus with a JN.1-like spike^31^. In turn, LF.7.2.1 (alias for JN.1.16.1.7.2.1) and LP.8.1 (alias for JN.1.11.1.1.1.3.8.1) are emerging variants with likely superior growth advantages over XEC^32^. In the early assessment and antigenic profiling of these strains, LF.7.2.1 was shown to be more immune evasive than XEC but with weaker ACE2 engagement^32^. In turn, LP.8.1 was shown to be as immune-evasive as XEC but with superior ACE2 engagement, which likely explains its improved transmission and rise in several countries^27, 32^. Both LP.8.1 and LF.7.2.1 are characterized by 8-9 spike differences from JN.1 including some RBD mutations (eg A475V and K444R in LF.7.2.1 and V445R in LP.8.1; **Fig. 4a**). Our risk assessment for these emerging variants against licensed vaccines, including mRNA-1273.167 and mRNA-1273.712, are shown in **Fig. 4b**. All of these variants are derived from the JN.1 S backbone, either via stepwise evolution or recombination, and are characterized by short-branches on the S phylogenetic tree (**Fig. 4c**) and in agreement with this, our predictive modeling suggests that both mRNA-1273.167 and mRNA-1273.712 are likely to cross-neutralize these emerging strains. Only LF.7.2.1 is predicted to be more immune-evasive, consistent with experimental observations, and has a risk score of ∼4 against mRNA-1273.167 and a risk score of ∼2.7 against mRNA-1273.712 vaccine but is likely to be outcompeted by fitter variants such as LP.8.1^32^.

**Fig. 4.**
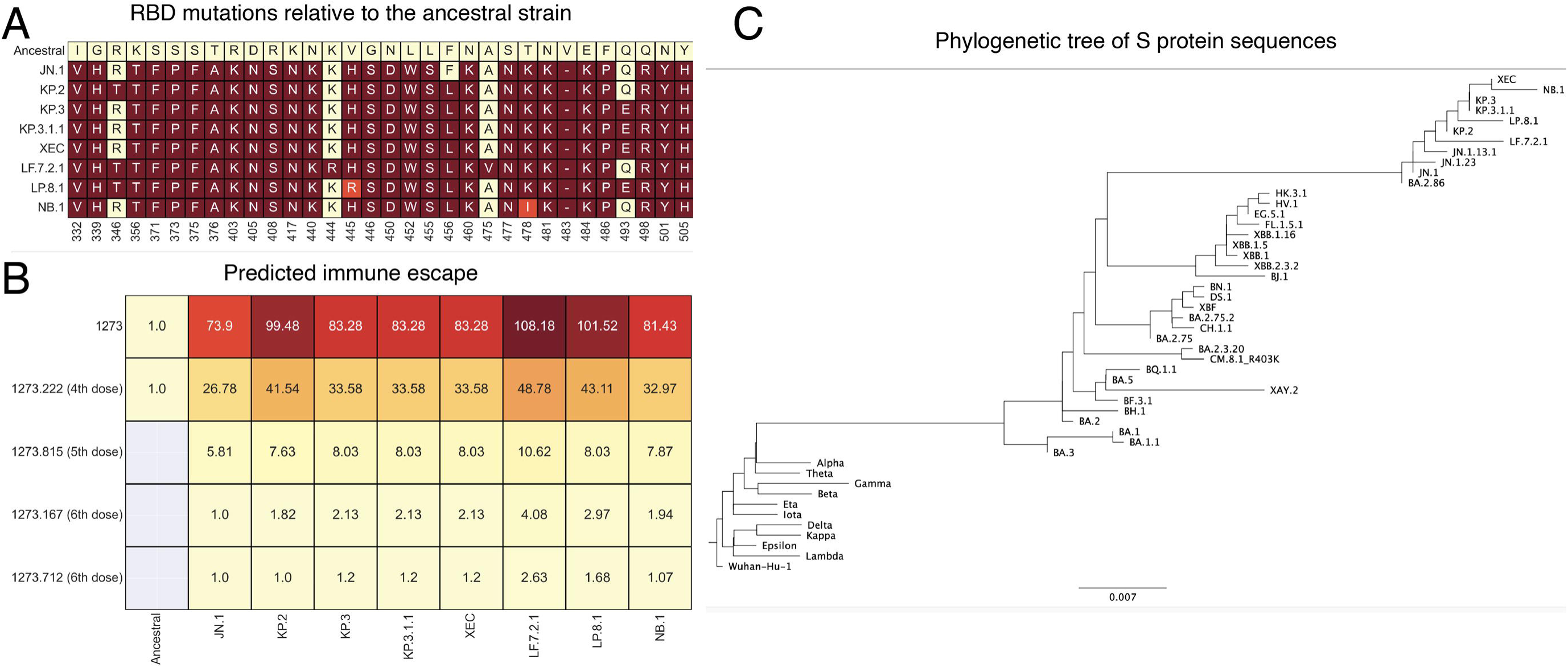
Antigenic grouping of emerging variants, predicted immune escape by vaccination history, and phylogeny of S variants. (A) S RBD differences among the emerging variants. The *x*-axis indicates the amino acid position relative to the ancestral (Wuhan-Hu-1) strain. Mutations are indicated in different colors. (B) The matrix shows predicted immune escape or potential of the emerging variants against recently approved and historical vaccines. The values inside the matrix are the predicted fold-drop in the ID_50_ neutralizing antibody GMTs on the vaccine strain. (C) A ML phylogeny describes the evolution of S variants. The tree is rooted in the branch leading to the ancestral (Wuhan-Hu-1) S sequence. The scale measures the average expected amino acid substitutions per site. Note that the branch lengths are influenced by recombination due to the presence of XBB, XEC, XBF, and XAY recombinant strains. GMT, geometric mean titer; ID_50_, 50% inhibitory dose; ML, maximum-likelihood; RBD, receptor-binding domain; S, spike glycoprotein.

### Clinical evaluations of the 2023-2024 vaccine composition support predictive risk modeling

The 2023-2024 vaccine composition (mRNA-1273.815, monovalent XBB.1.5) was evaluated as a fifth dose in the phase 2/3 Part J study (NCT04927065) in US adult participants (n = 50) who previously received a primary vaccination series of mRNA-1273, a third dose (booster) mRNA-1273, and a fourth dose (booster) of mRNA-1273.222. At Day 29 post-vaccination, mRNA-1273.815 induced robust neutralizing antibodies (nAbs) against the tested SARS-CoV-2 variants (**Fig. 5**). The nAb fold-increase from pre-booster levels were higher against the XBB.1.5, XBB.1.16, EG.5.1, BA.2.86, and JN.1 variants (10.4- to 19.0-fold) compared with the ancestral D614G, BA.4/BA.5, and BQ.1.1 variants (4.4- to 7.3-fold). The nAb titers at Day 29 after mRNA-1273.815 vaccination were lowest against the EG.5.1, BA.2.86, and JN.1, and were reduced by 2.5- to 5.8-fold compared with those elicited against the target variant XBB.1.5 (**Fig. 5**). Overall, the predicted escape based on the risk score was consistent with the experimentally assessed escape (nAb reduction) for all strains^11^.

**Fig. 5.**
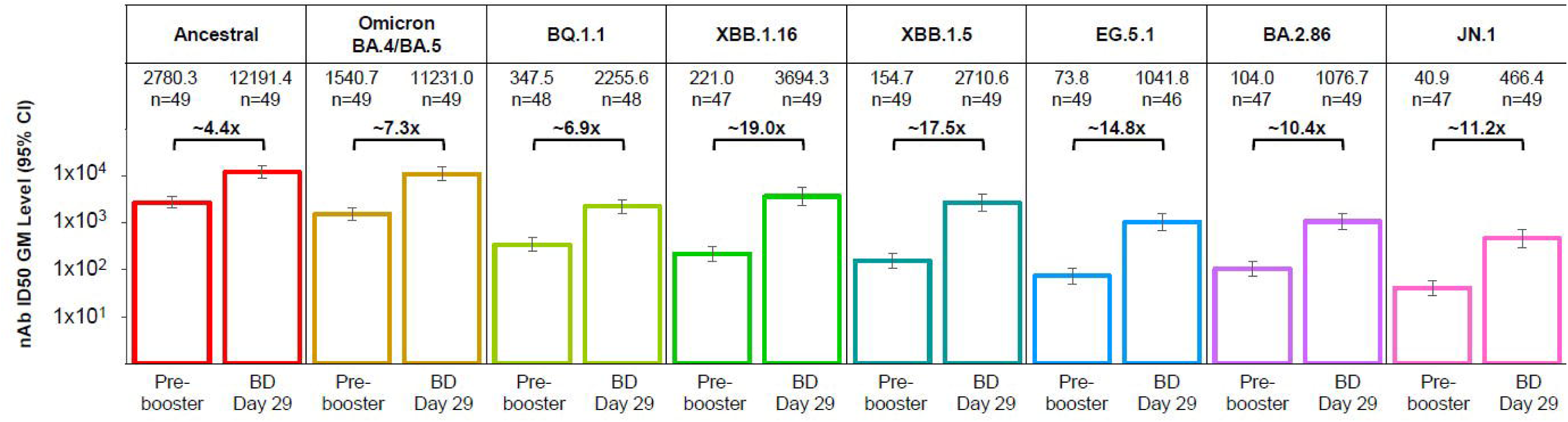
nAbs against SARS-CoV-2 variants pre-boost and at Day 29 after the mRNA-1273.815 booster. GMFR data are presented above the bars. BD, booster dose; GM, geometric mean; GMFR, geometric mean fold-rise; ID_50_, 50% inhibitory dose; nAb, neutralizing antibody. Adapted from Chalkias S, et al. *J Infect Dis.* 2024;230(2):e279-e286.^11^

## Discussion

Here we describe the utility of our risk calculator to predict the immune escape of emerging SARS-CoV-2 variants and to prioritize at-risk preparation of strains for annual and off-cycle vaccine updates, if needed. The risk calculator provides a quick assessment of potential immune escape of emerging variants and enables antigenic grouping of variants to predict cross-neutralization. Such data are informative both for internal decision-making and for at-risk preparation of vaccine constructs, if later requested by the global public health agencies. This is especially useful when there is co-circulation of multiple strains with comparable growth rates (eg, in case of incremental or stepwise convergent evolution of virus) but also helpful when more mutated strains emerge either via reverse zoonosis or chronic spillovers or recombination with some important differences, as discussed below.

While the risk calculator has great value in predicting the relative degree of immune escape exhibited by circulating strains against specific vaccines or immune backgrounds, some important limitations should be considered when interpreting the predicted risk scores.

First, the risk calculator is likely to be more accurate for variants that have evolved via stepwise evolution or antigenic drift by accumulating a small number of changes over time in a given backbone (eg, XBB and JN.1 subvariants from the parental strains included in the vaccines; **Fig. 4C**). This is because variants that primarily evolve via stepwise antigenic drift are more likely to be cross-neutralized by a closely matched vaccine strain (eg, JN.1 or KP.2 for the current season) and may not always prompt an urgent vaccine update. In turn, the calculator is likely to initially overestimate the immune escape of highly divergent strains that may have evolved via more dramatic evolutionary events, such as recombination or chronic spillovers. Such variants usually encode a large number of S and RBD mutations and manifest as long branches on phylogenetic trees (**Fig. 4C**), potentially representing saltation events that are difficult to predict or model and which could require urgent vaccine updates^33^. For example, the risk calculator initially overestimated the immune escape of the JN.1 variant against mRNA-1273.815 vaccine (predicted risk score of 15.73) when it first emerged, due to a lack of adequate model training data (eg, nAb titers for JN.1 vs. XBB.1.5 boosted individuals that would allow us to reinitialize the model parameters) and the observed genetic diversity of the new BA.2.86/JN.1 family of subvariants). In the case of such long branch variants that likely spillover from chronic infections, the S is likely to experience different or perhaps no immune pressure, resulting in the accumulation of a large number of mutations that may numerically suggest a greater degree of immune escape. Periodically updating, or retraining, to reinitialize the predictive model every time a new variant saltation becomes epidemiologically relevant allows the model to remain relevant as variant evolution progresses seasonally. Moreover, it remains important to continue to validate the predictions of the risk calculator experimentally and make adjustments, as needed.

Second, a large fraction of the global population is now likely exposed to multiple SARS-CoV-2 strains, either via repeated infections or vaccinations or both^34–36^. The order of infections and vaccinations also differs at the population level; infants are now primed only with the most recently approved vaccines, which may impact the specificity and target of the serum neutralization response^37^. Notably, the calculator is strictly trained on pandemic sera and antibodies (the majority targeting the RBD). This is a limitation, as NTD can have conformational effects on the RBD and may indirectly improve virus immune evasion, as highlighted by the recent success of KP.3.1.1 and XEC variants, which were both characterized by new NTD glycosylation sites^38^. Similarly, a recent study showed that infants primed/infected with only XBB strain had serum neutralization activity mostly targeted towards the NTD, in contrast to RBD-targeting neutralization activity in adults primed with the ancestral Wuhan-Hu-1 strain^37^. Such heterogeneities exist at the population level but are not modeled by the risk calculator^39^. Presently, the calculator does not distinguish between immunity acquired either via infection or vaccination or both. Similarly, the immune response to variants also involves additional components such as T-cell responses and host-related factors^40, 41^. We aim to implement these more complex features in future updates of the model.

Finally, >4,000 antibodies from previously published studies were used in the calculation of the Ab binding index and the training of the risk calculator^7–9^. This rich and comprehensive dataset naturally includes antibodies with different cross-reactivities (with the majority binding only to the ancestral Wuhan-Hu-1 strain); it is important to note that not all are nAbs (epitopes were determined via deep-mutation scanning) and that escape scores were aggregated over each site, thus ignoring the effects of different amino acid substitutions that may have different contributions to immune escape. Therefore, we are unable to model epistatic effects that are likely to manifest in different variant backbones. Similarly, the risk assessment model currently uses a training dataset for phase 2/3 Part H (mRNA-1273.222-boosted clinical sera) to make predictions about escape from the more recently approved vaccines. Ideally, it will be important to update the risk assessment model annually, coinciding with each new vaccine update or the emergence of a new saltation variant. However, logistical challenges that may occur, such as the availability of clinical sera and a sufficiently large number for pseudotyped variants to be included in model testing and training, have prohibited regular annual updates. We believe that even older models are appropriate or useful for making relative statements about immune escape, as there has been no fundamental change to the evolution of SARS-CoV-2 since its first emergence in 2019. The virus continues to evolve antigenically by acquiring new S mutations that can affect antibody binding and thereby neutralization; therefore, the inferred statistical relationship between antibody binding and neutralization loss is likely to remain robust through subsequent iterations of vaccines and variants. In other words, the actual numbers outputted by the risk calculator are likely to fluctuate season over season, especially when additional variants and antibodies are added to the training dataset of the model, but the relative patterns and relationships are likely to remain robust as highlighted by the remarkable consistency in the estimated models between two different clinical studies: the phase 2/3 Part H (**Figure 1**) and phase 1 study (**Fig. S1**) and the predicted risk scores versus experimentally determined escape scores in some published studies. Nevertheless, we continue to gather additional neutralization data using clinical sera from recent vaccinees and update the statistical models accordingly.

Despite these stated limitations, the risk calculator shows a strong correlation between antibody binding and predicted loss in neutralization, which is consistent across at least 2 clinical studies. As SARS-CoV-2 continues to evolve and spread, it remains important to prioritize strains rapidly for at-risk preparation, in case a vaccine update is needed or requested later by the health agencies. The risk calculator enables a quick quantitative assessment of immune escape for emerging subvariants to help manufacturers prioritize strains for experimental assessments from a much larger pool of virus genetic diversity and co-circulating variants. Risk scores generated by the model coupled with growth trends of variants provide valuable information about the ongoing adaptive evolution of SARS-CoV-2. We continue to perform annual updates to the risk calculator by integrating the latest pandemic sera from the sponsor’s ongoing clinical trials and deep-mutation scanning profiles of new antibody S complexes, as and when they become publicly available.

### Conclusions

By staying vigilant and proactive in our surveillance efforts, we can navigate the complex and evolving landscape of SARS-CoV-2 variants, adapt and optimize our strategies accordingly, and ultimately safeguard the health and well-being of populations worldwide. Here, we describe our response to the ongoing adaptive evolution of SARS-CoV-2 based on routine and active genomic surveillance combined with bioinformatics-based risk assessment. Our risk calculator allows us to identify candidate vaccine strains within a condensed time window to support seasonal updates, if later requested by the health agencies. We continue to monitor SARS-CoV-2 evolution and assess and report on the impact of new variants against the fall 2024-2025 JN.1 and KP.2 vaccine formulations, including the assessment of variants that become globally dominant and those variants that show potential for immune escape from the current season’s vaccine composition.

## Data Availability

Access to participant-level data presented in this article and supporting clinical documents with external researchers who provide methodologically sound scientific proposals will be available upon reasonable request for products or indications that have been approved by regulators in the relevant markets and subject to review from 24 months after study completion. Such requests can be made to Moderna, Inc., 200 Technology Square, Cambridge, MA 02139 < > and to the corresponding author at < >. A materials transfer and/or data access agreement with the sponsor will be required for accessing shared data. All other relevant data are presented in the paper. The protocol is available online at ClinicalTrials.gov: NCT04927065. The example genomic sequence data and metadata for variants reported in this study can be accessed via EPI_SET_250127qx. To view the contributors of each individual sequence with details such as accession number, virus name, collection date, originating lab and submitting lab and the list of authors, visit 10.55876/gis8.250127qx.

## Acknowledgements

We gratefully acknowledge all data contributors, ie, the authors and their originating laboratories responsible for obtaining the specimens, and their submitting laboratories for generating the genetic sequence and metadata and sharing via the GISAID Initiative, on which some of this research is based. We thank Jesse Bloom and Yunlong Cao for providing critical data on antibody S epitopes and escape profiles that were used in the development of the risk calculator and for useful discussions on the manuscript. We are grateful to Frances Priddy, Spyros Chalkias, and Bethany Girard for their significant contributions to the clinical trial data referenced in this manuscript and Gwo-Yu Chuang, Nihal Korkmaz, Yen-Ting Lai, and Swan Tan (all Moderna, Inc.) for useful discussions and contributions that led to the development of the current model. We also thank Honghong Zhou and Jing Chen (both statisticians at Moderna Inc.) for a consultation and review of the statistical model developed and described in this study. Writing and/or editorial assistance was provided by Rachel Lesson (Moderna, Inc.) and MEDiSTRAVA in accordance with Good Publication Practice (GPP 2022) guidelines, funded Moderna, Inc., and under the direction of the authors. This study was funded by Moderna, Inc.

## Author Contributions

AN, LA, AC, GS-J, DE contributed to study concept and design. AN, DL, LA, DMB, TS, GS-J, DE contributed to data collection. AN, DL, LA, DMB, TS, YB, GS-J, DE contributed to data analysis/interpretation. AN, DL, LA, KW, YB, AC, GS-J, DE made intellectual contributions and contributed to writing and review of the manuscript. All authors approved the final version of the manuscript.

