## Supplementary Figure 1 for "Predictive Modeling of Immune Escape and Antigenic Grouping of SARS-CoV-2 Variants"

**Fig. S1**

Statistical relationship between the antibody binding index ($x$-axis) and serum neutralization for SARS-CoV-2 variants evaluated against the clinical sera from the mRNA-1273 phase 1 study ($y$-axis). The estimated regression function is listed, along with the statistical test for significance

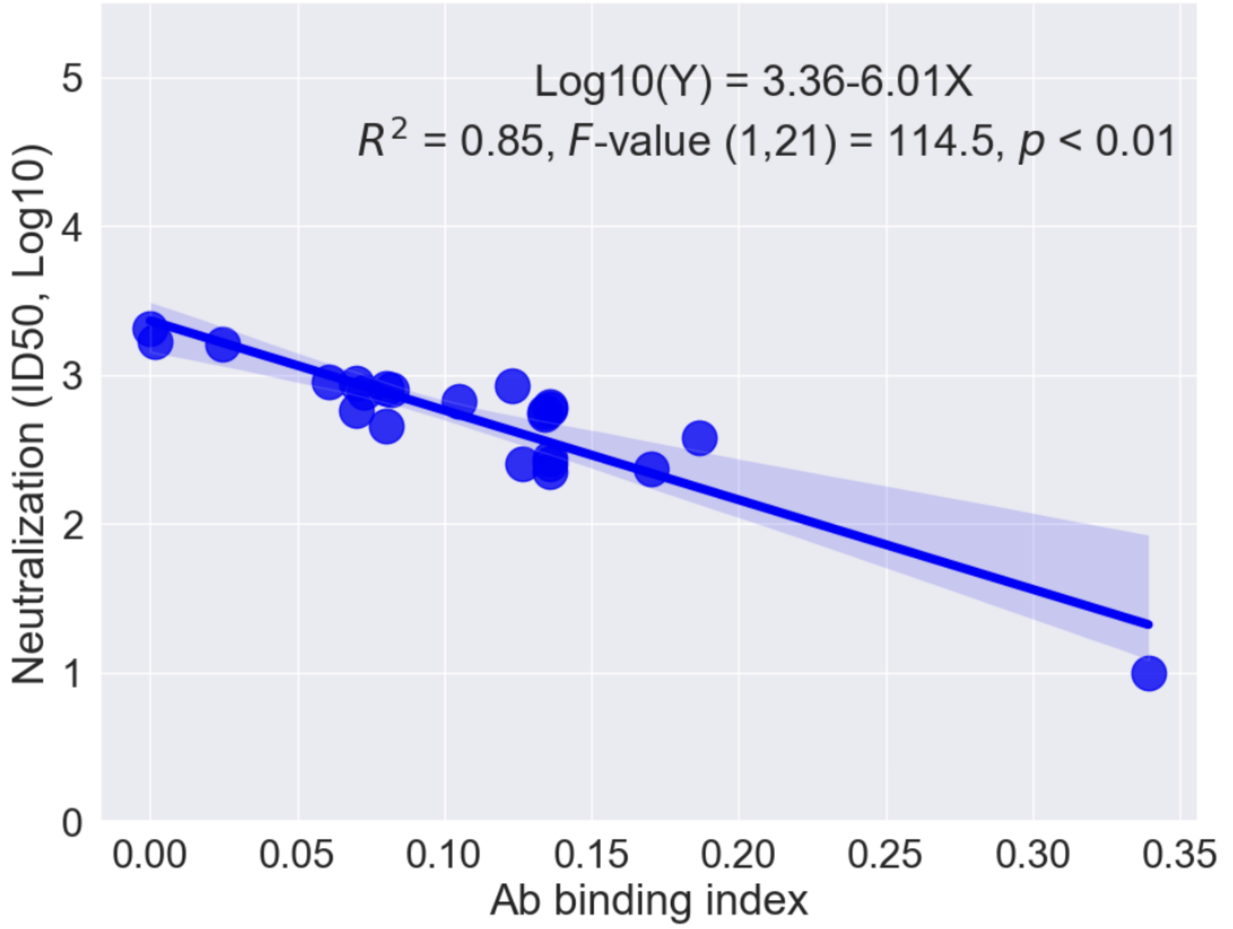


Ab, antibody; $ID_{50}$, 50% inhibitory dose.
